# Catalytic cleavage of HEAT and subsequent covalent binding of the tetralone moiety by the SARS-CoV-2 main protease

**DOI:** 10.1101/2020.05.02.043554

**Authors:** Sebastian Günther, Patrick Y. A. Reinke, Dominik Oberthuer, Oleksandr Yefanov, Helen Ginn, Susanne Meier, Thomas J. Lane, Kristina Lorenzen, Luca Gelisio, Wolfgang Brehm, Illona Dunkel, Martin Domaracky, Sofiane Saouane, Julia Lieske, Christiane Ehrt, Faisal Koua, Alexandra Tolstikova, Thomas A. White, Michael Groessler, Holger Fleckenstein, Fabian Trost, Marina Galchenkova, Yaroslav Gevorkov, Chufeng Li, Salah Awel, Ariana Peck, P. Lourdu Xavier, Miriam Barthelmess, Frank Schlünzen, Nadine Werner, Hina Andaleeb, Najeeb Ullah, Sven Falke, Bruno Alves Franca, Martin Schwinzer, Hévila Brognaro, Brandon Seychell, Henry Gieseler, Diogo Melo, Joanna I. Zaitseva-Kinneberg, Brenna Norton-Baker, Juraj Knoska, Gisel Esperanza, Aida Rahmani Mashhour, Filip Guicking, Vincent Hennicke, Pontus Fischer, Cromarte Rogers, Diana C. F. Monteiro, Johanna Hakanpää, Jan Meyer, Heshmat Noei, Phil Gribbon, Bernhard Ellinger, Maria Kuzikov, Markus Wolf, Linlin Zhang, Xinyuanyuan Sun, Jonathan Pletzer-Zelgert, Jan Wollenhaupt, Christian Feiler, Manfred Weiss, Eike-Christian Schulz, Pedram Mehrabi, Christina Schmidt, Robin Schubert, Huijong Han, Boris Krichel, Yaiza Fernández-García, Beatriz Escudero-Pérez, Stephan Günther, Dusan Turk, Charlotte Uetrecht, Tobias Beck, Henning Tidow, Ashwin Chari, Andrea Zaliani, Matthias Rarey, Russell Cox, Rolf Hilgenfeld, Henry N. Chapman, Arwen R. Pearson, Christian Betzel, Alke Meents

## Abstract

Here we present the crystal structure of SARS-CoV-2 main protease (M^pro^) covalently bound to 2-methyl-1-tetralone. This complex was obtained by co-crystallization of M^pro^ with HEAT (2-(((4-hydroxyphenethyl)amino)methyl)-3,4-dihydronaphthalen-1(2H)-one) in the framework of a large X-ray crystallographic screening project of M^pro^ against a drug repurposing library, consisting of 5632 approved drugs or compounds in clinical phase trials. Further investigations showed that HEAT is cleaved by M^pro^ in an E1cB-like reaction mechanism into 2-methylene-1-tetralone and tyramine. The catalytic Cys145 subsequently binds covalently in a Michael addition to the methylene carbon atom of 2-methylene-1-tetralone. According to this postulated model HEAT is acting in a pro-drug-like fashion. It is metabolized by M^pro^, followed by covalent binding of one metabolite to the active site. The structure of the covalent adduct elucidated in this study opens up a new path for developing non-peptidic inhibitors.

## Introduction

The pandemic disease caused by the coronavirus SARS-CoV-2 is an international health challenge and there is an immediate need for the identification of drugs for treatment of SARS-CoV-2 infections. Infection of host cells critically depends on several molecular factors from both the host and virus (*1, 2*). Coronaviruses are RNA-viruses with a genome of around 30 kbp. The viral gene essential for the replication of the virus is expressed as polyprotein that must be broken into functional subunits for replication and transcription activity (*1*). This proteolytic cleavage is primarily accomplished by the main protease (M^pro^), also known as the 3C-like protease 3CL^pro^. M^pro^ cleaves the viral polyprotein at eleven sites. The core cleavage motif is Leu-Gln⇓(Ser/Ala/Gly) (*1*). M^pro^ possesses a chymotrypsin-like fold amended by a C-terminal helical domain and harbors a catalytic dyad comprising Cys145 and His41 (*1*). The active site is located in a cleft between the two N-terminal domains of the three domain structure of the monomer, while the C-terminal helical domain is involved in regulation of dimerization of the enzyme with a dimerization constant of 2.5 µM (*1*). Due to its central involvement in virus replication M^pro^ is well established as a prime target for antiviral compounds that can tackle coronavirus infections (*3*). Indeed first studies confirm the potential of targeting M^pro^ for inhibition of virus replication (*1, 2*).

In order to identify further potential drug candidates with an inhibitory effect on M^pro^ we screened the “Fraunhofer IME Repurposing Collection” (*4*), consisting of 5632 individual compounds against M^pro^ by means of high-throughput X-ray crystallography. This compound collection is based on the BROAD institute repurposing library (*5*). In contrast to crystallographic screening campaigns with small molecules (“fragment screening”) (*6*), our library contains compounds that are chemically more complex and of larger molecular mass. As M^pro^ crystals possess a low solvent content of only 38% (*1*) with narrow solvent channels that could impede larger molecules from diffusing into the active site in soaking experiments, we co-crystallized M^pro^ with the individual compounds instead of soaking pre-grown M^pro^ crystals in compound-containing solutions. Crystals typically appeared within 2 days and were then submitted to single crystal X-ray structure determination (see Methods). From the resulting X-ray structures, several compounds were identified in complex with M^pro^. Among these was BE-2254 (HEAT, 2-(((4-hydroxyphenethyl)amino)methyl)-3,4-dihydronaphthalen-1(2H)-one), where a well resolved electron-density was observed in the active site of the M^pro^ crystal structure. This electron-density can be well modeled as 2-methyl-1-tetralone (METT) covalently bound to the sulfur atom of Cys145, thereby blocking access to the active site of M^pro^ and occupying the P1 site of the protease. 2-Methyl-1-tetralone is potentially a breakdown product of the parent compound HEAT included in the co-crystallization experiment.

HEAT was first synthesized by Hansen at Beiersdorf AG (*7*). It exists in an equilibrium between two enantiomers between which it transitions via keto-enol tautomerism (*8*). Investigations by several groups found it to be a competitive α-adrenoreceptor antagonist *in vitro* and *in vivo* with lower affinity to the α-subtype and residual affinity to dopaminergic and serotonergic receptors (*9*).

Here we present the crystal structure of M^pro^ with 2-methyl-1-tetralone (METT) covalently bound to the active site to 1.8 Å resolution. To further support this observation and to understand the underlying mechanism of HEAT fragmentation by M^pro^ and its subsequent covalent binding to the active site, we conducted complementary native and small molecule mass spectrometry and computational docking studies. On the basis of these results we propose a reaction mechanism and provide an outlook for the application of HEAT as a starting point for potential drug development to treat SARS-CoV-2 infections.

## Results

The overall structure of the SARS-CoV-2 main protease (M^pro^) with bound METT as obtained in our X-ray diffraction experiment is shown in figure 1 (table 1). It possesses the same overall structural features as previously described (*1, 2*). A closer view of the active site reveals clear unoccupied electron-density for a small molecule covalently bound to the catalytic Cys145 which could be well modeled as METT (figure 1B). The chemical structures of the parent compound HEAT and the observed cleavage product METT bound to M^Pro^ are illustrated in figure 1C and D, respectively, clearly showing the relationship between the two molecules. This observation suggests METT to be a breakdown product of HEAT, with tyramine (TY) as the complementary product (figure 1E).

**Table 1.** Data collection and refinement statistics (molecular replacement)

|  | M<sup>pro</sup>/2-methyl-1-tetralone |
| --- | --- |
| **PDB** | 6YNQ |
| **Data collection** |  |
| Space group | C2 |
| Cell dimensions |  |
| *a*, *b*, *c* (Å) | 113.1, 53.2, 44.7 |
| $\alpha$, $\beta$, $\gamma$(°) | 90, 103, 90 |
| Resolution (Å) | 27.29 - 1.8 (1.864 - 1.8) |
| *R*<sub>sym</sub> or *R*<sub>merge</sub> | 0.113 (1.853) |
| *CC*1/2 | 0.997 (0.435) |
| *mean I* / σ*I* | 8.1 (0.9) |
| Completeness (%) | 97.37 (95.85) |
| Redundancy | 3.8 (3.9) |
| **Refinement** |  |
| Resolution (Å) | 27.29 - 1.8 |
| No. reflections | 23506 (2309) |
| *R*<sub>work</sub> | 0.1821 (0.3661) |
| *R*<sub>free</sub> | 0.2285 (0.4468) |
| No. of non-hydrogen atoms | 2839 |
| Protein | 2528 |
| Ligand/ion | 17 |
| Solvent | 294 |
| *B*-factors (Å<sup>2</sup>) |  |
| Protein | 33.01 |
| Ligand/ion | 44.01 |
| Solvent | 38.3 |
| R.m.s. deviations |  |
| Bond lengths (Å) | 0.011 |
| Bond angles (°) | 0.97 |

\*Values in parentheses are for highest-resolution shell.
Data was collected from a single crystal

**Figure 1:**
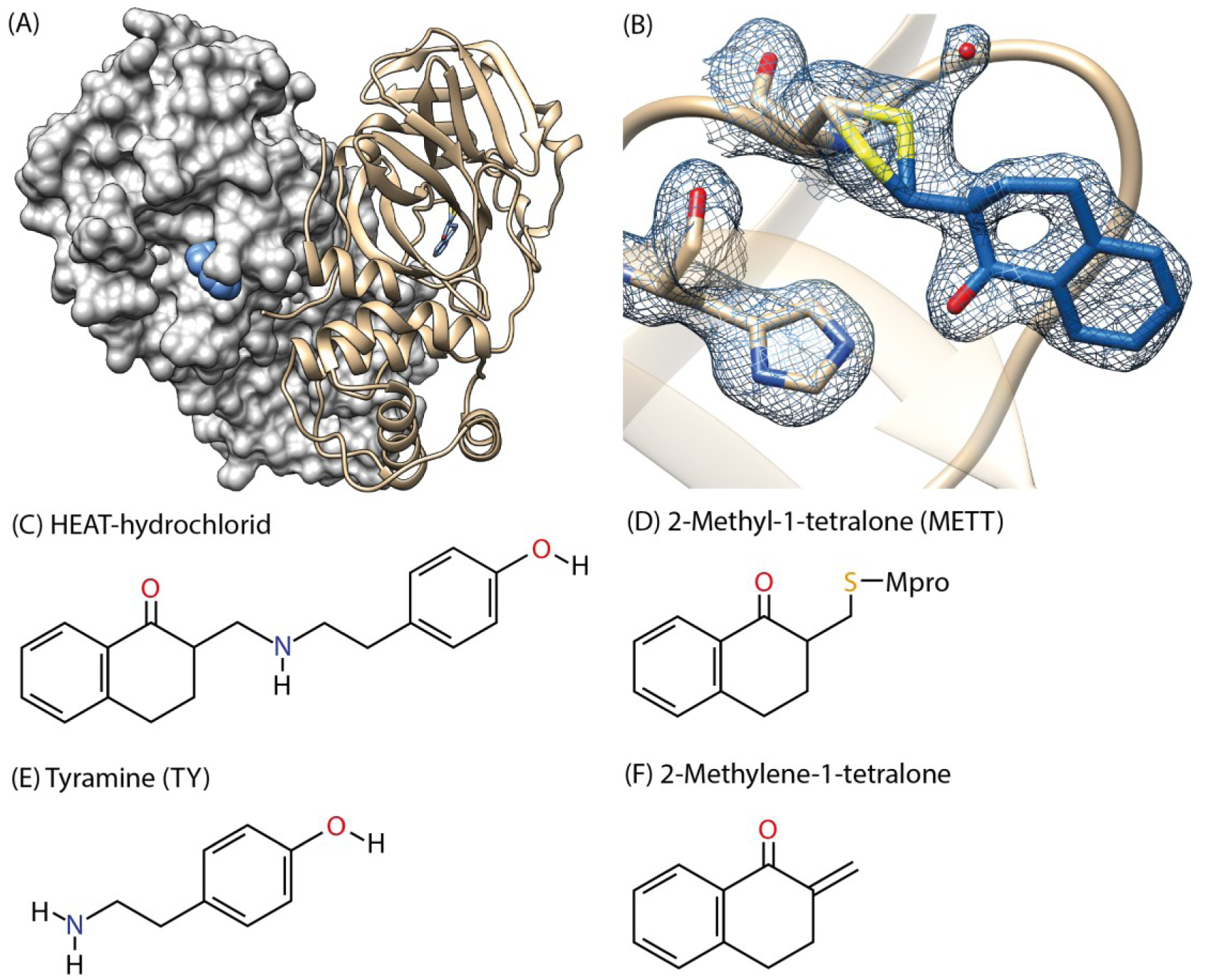
X-ray structure of M^pro^ with covalently bound 2-methyl-1-tetralone. (A) The biologically active form of M^pro^ is a homodimer found in the crystal structure. Here one monomer is depicted as surface representation, while the second monomer is shown as cartoon model. The ligand is highlighted as sphere and stick model (blue). (B) A simulated annealing omit 2Fo-Fc-map of the active site of M^pro^ reveals clear electron-density for a ligand covalently bound to Cys145 that can be modeled as METT. Coordinates of the ligand were removed before calculating the map (1 rmsd and carved at 1.4 Å around the atoms). Diffraction data together with coordinates were deposited in the Protein Data Bank with PDB accession code 6YNQ. (C-F) Chemical structures of the parent compound HEAT (C), 2-methyl-1-tetralone (METT) bound to M^pro^ (D), the proposed degradation products tyramine (TY, E) and the potential reaction intermediate 2-methylene-1-tetralone (F).

To confirm the cleavage of HEAT by M^pro^ and subsequent covalent binding of METT to M^pro^, native mass spectrometry (MS) experiments were performed (*10*). In a control experiment without the addition of HEAT we observe a high intensity peak envelope corresponding to the expected mass of the M^pro^ dimer (67,599 +/-5 Da, charge states +14 to +16). The monomer is also visible with lower intensity (33,795 +/-2 Da, charge states +9 to +11) (figure 2A). Next M^pro^ was incubated with HEAT at a molar ratio of 1:10 (M^pro^:HEAT) overnight. After incubation of M^pro^ with HEAT two additional peak envelopes can be observed in the high mass range (figure 2B). Their assigned masses correspond to the M^pro^ dimer alone or plus one and two METT molecules (67,593/67,774/67,955 Da). The corresponding stoichiometry of M^pro^ dimer without or either one or two bound METT was approximately 4:5:1. Additionally peaks from monomeric M^pro^ bound to one METT molecule and unmodified M^pro^ are also observed. The expected ion of intact HEAT with 296 Da (MH^+^) is observed in the low mass range (suppl. fig. 1). The reaction product METT is not observed as it is too small to be detected with the instrument settings used for these measurements.

**Figure 2:**
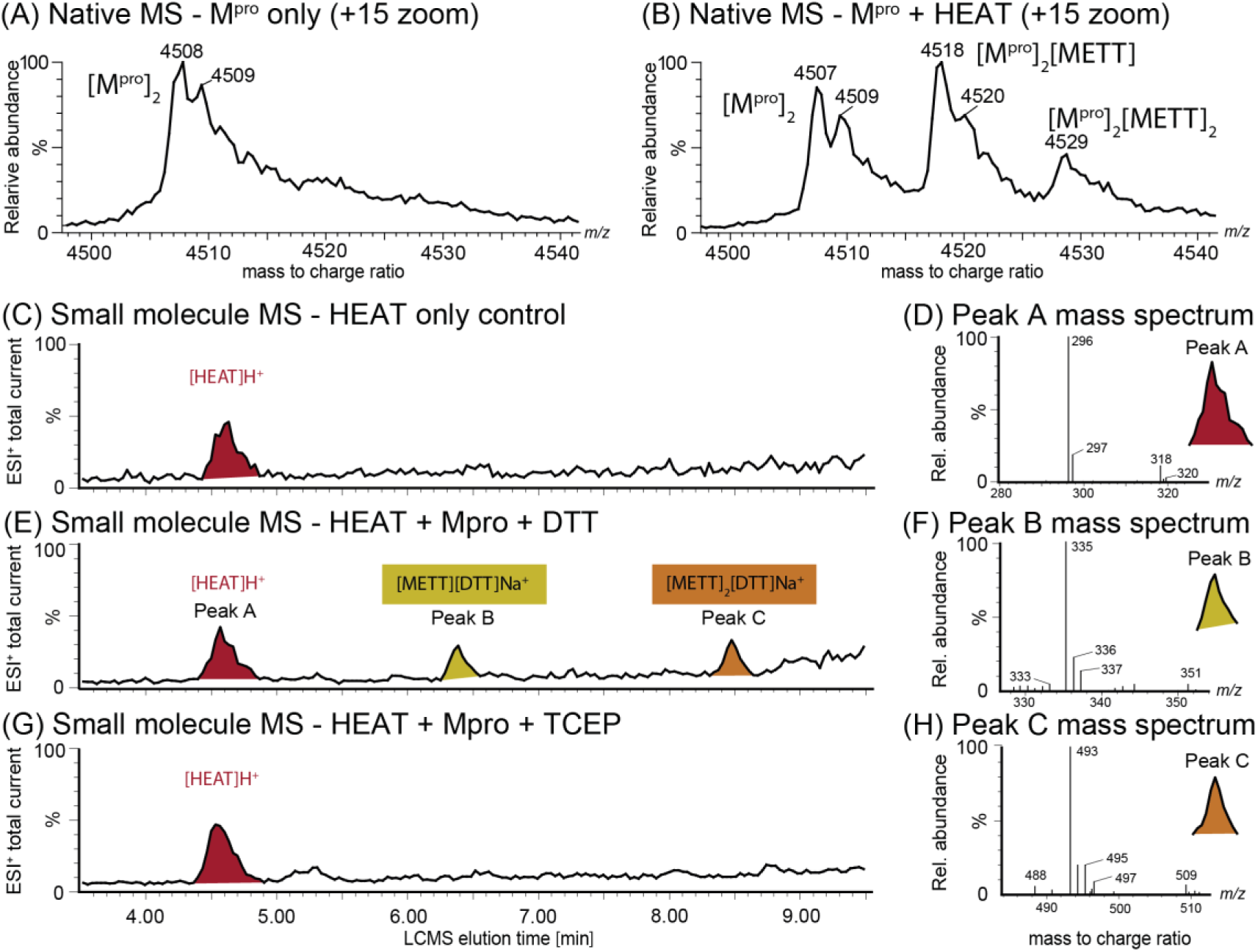
MS analysis of the educts and products of the reaction of HEAT with M^pro^. (A) native mass spectrum of native M^pro^ indicating the homodimeric state. (B) mass spectrum of M^pro^ after overnight incubation with HEAT leading to additional peaks with higher masses. (C-H) HPLC elution profile (ESI^+^, Total Ion Current) of the reaction products of HEAT alone (C) and with M^pro^ in the presence of DTT (E) or TCEP (G). The mass spectra corresponding to the elution peaks are shown in D ([HEAT]H^+^), F ([METT][DTT]Na^+^) and H ([METT]_2_[DTT]Na^+^).

To probe the chemical interaction between M^pro^ and HEAT, CID-MS/MS was performed by separating a molecular ion precursor (+15, dimer) in the quadrupole of the mass spectrometer and triggering its dissociation in a collision cell (see Methods). The dimer dissociated into low- and high-charge monomeric molecular ions. METT was observed to still be bound to the high-charge ions (suppl. figure 1). The ratio of the dissociated monomer in the unbound to bound state was roughly 5:4. The binding of the reactive group to the dissociated molecule indicates a strong interaction between M^pro^ and METT, as would be expected for a covalent bond. Furthermore, only one METT per monomer is detected which suggests that the interaction is site-specific.

In addition, the small-molecule products from the reaction of HEAT with M^pro^ were analyzed by liquid chromatography (LC)-MS in the range of 100 to 1000 Da (figure 2, see Methods). Pure HEAT elutes at 4.5 min and displays major ions at 296 Da (MH^+^) and 318 Da (MNa^+^). After overnight incubation (12 hours) at room temperature of an excess of HEAT with M^pro^, two new LC-MS peaks were observed at 6.4 and 8.5 min. Masses of 335.3 and 493.3 Da appear in the corresponding MS-ES^+^ spectrum, which could not be attributed to either of the expected reaction products METT (183 Da) or TY (137 Da). These observed masses correspond to the two possible adducts of METT and DTT (154 Da), which was used as reducing agent in the protein buffer. The observed signals correspond to MNa^+^ ions of METT bound to one or both thiol groups of DTT ([METT]_2_[DTT]Na^+^ or [METT] [DTT]Na^+^, respectively). The reaction product TY was observed in the MS-ES^-^ spectrum (137 Da, supp. figure 2) in the presence of M^pro^. To confirm the assignment of the METT and DTT adducts we substituted DTT with tris(2-carboxyethyl)phosphine (TCEP) as reducing agent. In the absence of DTT no additional LC-MS signals besides HEAT were observed, and no masses corresponding to the METT-DTT adducts could be detected. This supports the presence of the proposed METT intermediate (figure 1F) in the HEAT cleavage mechanism, which can be intercepted by the strong nucleophilic thiol group of DTT. In contrast, TCEP is much less nucleophilic and hence does not react with the intermediate of the HEAT cleavage.

Understanding of the underlying reaction mechanism requires knowledge about the initial binding of HEAT to M^pro^ which was not observed in the X-ray diffraction experiment. To obtain information on the initial binding state of HEAT to M^pro^, computational docking studies were employed using JAMDA and HYDE (*11*) (figure 3). We visually inspected multiple proposed non-covalent binding modes which might help to reveal the possible reaction mechanism. The keto form of HEAT (figure 3A) is found to occupy pockets P1 and P2 with a carbon-sulfur distance of 3.4 Å. HEAT forms a hydrogen bond to His163 in the active site via the carbonyl oxygen atom, a weak hydrogen bond to Cys145 via its secondary amine, and a T-stacking interaction to His41. The enol form (figure 3B) seems to be more stably bound to the active site, forming not only the two aforementioned hydrogen bonds, but also a hydrogen bond to Cys44 via its phenol moiety. Additionally, a pi-pi stacking interaction between His41 and the phenol moiety can be observed in this predicted binding mode. Moreover, the carbon-sulfur distance of 3.5 Å for the enol form of HEAT to Cys145 is well within the acceptable range for covalent bond formation. This non-covalent binding mode corresponds well to the experimentally determined binding mode of METT which occupies the S1 pocket (figure 3C). Additionally, the leaving group of the molecule is situated in the S2 pocket and is engaged in a weak pi-pi stacking interaction with His41 and a weak hydrogen bond to Cys44. We hypothesize that HEAT initially binds non-covalently to the protease and is converted to the corresponding enol form prior to the formation of the α,β-unsaturated ketone. This process might be catalyzed by residues His163 and Cys145, which are engaged in hydrogen bonds to the product METT as well as the proposed non-covalently bound keto- and enol-form of HEAT. Due to the covalent binding to the active-site Cys145, we expected an inhibitory effect on the enzyme function and thus virus replication.

**Figure 3:**
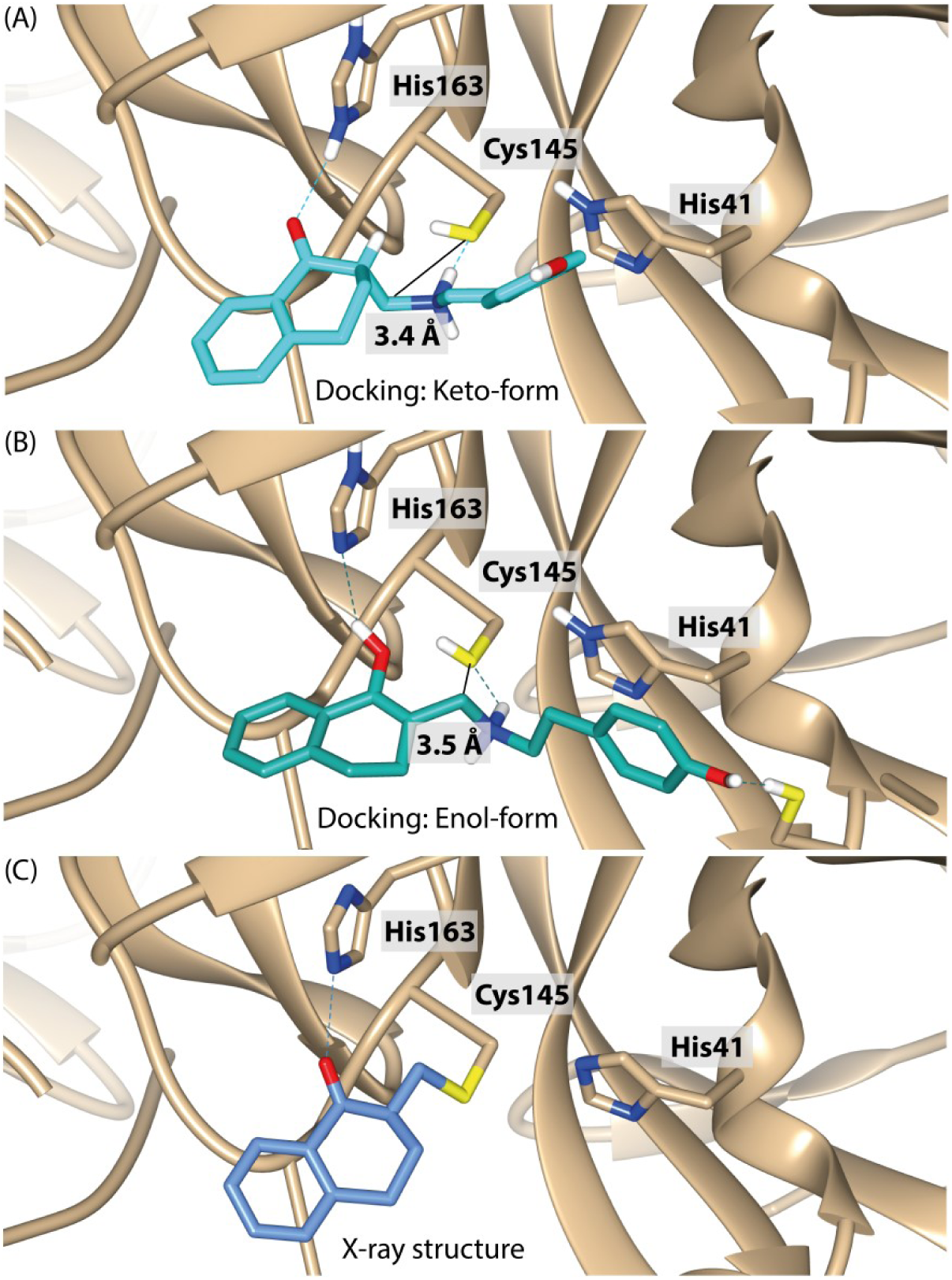
Proposed non-covalent binding modes of HEAT in comparison to the experimentally determined covalent binding mode. Docking studies were used to obtain information about the possible binding modes of M^pro^ to the keto-form (A) and enol-form (B) of HEAT. Hydrogen bonds are depicted as dotted lines and the carbon-sulfur distance is highlighted in black. For comparison the experimental binding mode of 2-methyl-1-tetralone as observed in the X-ray structure is shown (C).

## Discussion

Combining all experimental and computational results allows us to propose a plausible, albeit highly unusual, reaction mechanism. M^pro^ is a cysteine protease possessing the conserved reactive center dyad Cys145 and His41, which together catalyze the sequence-specific hydrolysis of peptide bonds, vital for SARS-CoV-2 replication. We propose a similar mechanism for the cleavage of the drug HEAT, leading to a suicide reaction of the protease, which terminates in a covalent thioether bond between Cys145 and METT (figure 4). As initiation the tetralone carbonyl is activated by His163, forming an enol. The thiolate of Cys145 could serve as base, removing the hydrogen of the β-carbon and forming the enol. After enol-oxygen deprotonation by His163, an unimolecular conjugate base elimination (E1cB)-like mechanism expels the protonated tyramine amine, forming 2-methylen-1-tetralone in the active site. In a subsequent step this highly reactive intermediate reacts in a Michael addition, where the thiolate of Cys145 attacks the α,β-unsaturated 2-methylen-1-tetralone, forming a thioether bond resulting in covalently bound METT. This final reaction state was confirmed by X-ray crystallography to a resolution of 1.8 Å. As evident in our LC-MS analysis some METT can react with DTT and it is not yet clear whether this occurs within the active site or spontaneously in solution.

**Figure 4:**
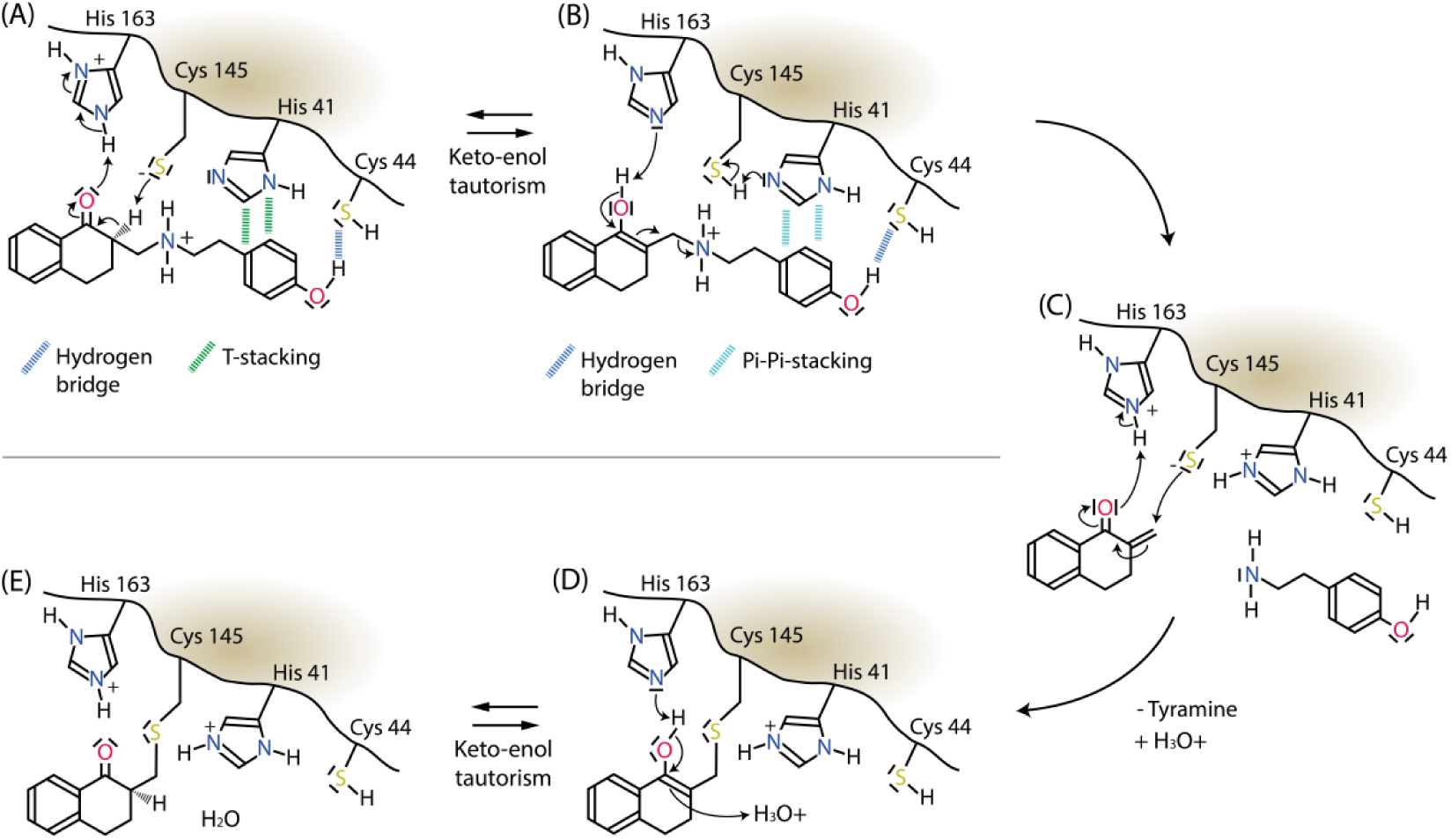
Proposed reaction mechanism of HEAT with M^pro^. HEAT exists in equilibrium between a keto- (A) and enol- (B) form via tautomerism. The catalytic center bound enol form of HEAT eliminates the tyramine group, leaving 2-methylene-1-tetralone in the active site (C). This is attacked by the thiol of Cys145, forming the observed covalent thioether bond (D). This enol is again in equilibrium with the keto-form, observed in the X-ray structure (E).

The current pandemic of SARS-CoV-2 and also the earlier epidemics caused by SARS-CoV and MERS-CoV revealed the risk posed by coronaviruses and stressed the need for pan-coronavirus antiviral drugs to be better prepared for future outbreaks. M^pro^ is highly conserved across coronaviruses (*12*) (suppl. figure 3). The active site residues His41 and Cys145 are strictly conserved. Furthermore, in our structure METT binding is stabilized by a hydrogen bond to His163 as well as by van der Waals interactions with the backbone of the oxyanion hole formed by residues Leu141 and Asn142 and neighboring Phe140. These residues are all essential for the protease activity of M^pro^ and thus are conserved as well. Further experiments to investigate the influence of HEAT on the activity of other coronaviruses need to be conducted to confirm the potential of developing HEAT as a pan-coronavirus inhibitor.

Currently most structurally well described SARS-CoV-2 M^pro^ inhibitors are rationally designed peptidomimetic or peptide-based inhibitors (*1, 2, 13, 14*). However, some structures of non-peptidic inhibitors were recently reported. For example Baicalein is a natural product derived from traditional Chinese medicine (*15*). It is non-covalently bound in the active site of M^pro^. In contrast the antineoplastic drug carmofur is converted by M^pro^ and its lipophilic hexamethylcarbamate moiety remains covalently bound to Cys145 (*16*), likely through a mechanism reminiscent of hydrolyzing a peptide substrate. Chemically the mechanism of reaction of carmofur with M^pro^ is different from the one we observe for HEAT.

In conclusion we have discovered that the alpha adrenergic antagonist HEAT is metabolized by and covalently bound to the active site of SARS-CoV-2 M^pro^. Further evaluation of the antiviral activity of HEAT is required, as in a first high throughput screen of the same repurposing library it did not show any significant antiviral activity (*4*). In spite of this, the complex between METT and M^pro^ can be considered a starting point for future drug design efforts. Chemical modifications of the molecule might help to endow it with pharmacologically relevant inhibitory activity and to develop HEAT towards a viable drug option.

The obtained results also strongly highlight that drug discovery investigations to treat SARS-CoV-2 should not only focus on traditional and conservative approaches aiming to identify substrate-analogue inhibitors, but should also include the identification and discovery of pro-drug compounds that could act as suicide drugs (*17, 18*). This approach is based on the simple but effective idea of a classical pro-drug, which needs to be metabolized first before becoming toxic for the infecting agent, in this case the virus SARS-Cov-2 via deactivation of M^pro^.

In this context and according to our postulated model, HEAT can be considered as a pro-drug as well as suicide drug which binds first to the active site of M^pro^, where it is cleaved in an E1cB-like reaction mechanism into 2-methylene-1-tetralone and tyramine. The Cys145 sulfur subsequently binds covalently in a Michael addition to the methylene carbon atom of 2-methylene-1-tetralone thereby blocking the active site and inactivating M^pro^.

## Methods

### M^pro^ crystallization

The protein was overexpressed in *E. coli* and purified for subsequent crystallization according to previously published protocols (*1*). Co-crystallization with the compounds was achieved by equilibrating a 6.25 mg/ml protein solution in 20 mM HEPES buffer (pH 7.8) containing 1 mM DTT, 1 mM EDTA, and 150 mM NaCl against a reservoir solution of 100 mM MIB, pH 7.5, containing 25% w/w PEG 1500 and 5% *v/v* DMSO. Prior to crystallization compound solutions in DMSO were dried onto the wells of SwissCI 96-well plates. To obtain well-diffracting crystals in a reproducible way seeding was applied for crystal growth. Crystals appeared within a few hours and reached their final size after 2 - 3 days. Crystals were manually harvested and flash-frozen in liquid nitrogen for subsequent X-ray diffraction data collection.

### X-ray data collection and processing

X-ray data were collected at beamline P11 at the synchrotron radiation facility PETRA III at DESY in Hamburg(*19*). Data were collected at an X-ray energy of 12 keV. Using the P11 sample changing robot data collection time was 3 minutes for a full 200 degree rotation dataset, including sample exchange. Data analysis was carried out using XDS (*20*), COOT (*21*), PHENIX (*22*), and PanDDA (*23*), in addition to some bespoke software tools. Chemical restraints for the bound compound were generated from SMILES using phenix.elbow (*24*) and compared with results from acedrg (*25*) and grade (*26*). The fully refined model and structure factors were deposited in the PDB with accession code 6YNQ. The distribution of protein residues over the Ramachandran plot was 97.04/2.63/0.33% (favored/allowed/outliers).

### Mass-spectrometric analysis of compound binding

M^pro^ samples were prepared for native MS measurements by buffer-exchange into ESI compatible solutions. M^pro^ was buffer exchanged into 300 mM NH_4_OAc, pH 7.5 by five cycles of centrifugal gel filtration (Vivaspin columns, 30.000 MWCO, Sartorius). M^pro^ (250 µM, 300 mM NH_4_OAc, pH 7.5) was incubated with HEAT (2.5 mM in DMSO) for 20 h and diluted 1:20.

Nano ESI capillaries were pulled in-house from borosilicate capillaries (1.2 mm outer diameter, 0.68 mm inner diameter, filament, World Precision Instruments) with a micropipette puller (P-1000, Sutter instruments) using a squared box filament (2.5 × 2.5 mm^2^, Sutter Instruments) in a two-step program. Subsequently capillaries were gold-coated using a sputter coater (Q150R, Quorum Technologies) with 40 mA, 200 s, tooling factor 2.3 and end bleed vacuum of 8×10^−2^ mbar argon. Native MS was performed using an electrospray quadrupole time-of-flight (ESI-Q-TOF) instrument (Q-TOF2, Micromass/Waters, MS Vision) modified for higher masses (*27*). Samples were ionized in positive ion mode with voltages applied at the capillary of 1300 V and at the cone of 130 V. The pressure in the source region was kept at 10 mbar throughout all native MS experiments. For desolvation and dissociation, the pressure in the collision cell was adjusted to 1.8×10^−2^ mbar argon. Native-like spectra were obtained at an accelerating voltage of 30-100 V, while for collision-induced dissociation these voltages were increased to 150-170 V. In ESI-MS overview spectra, the quadrupole profile was 1-10,000 m/z. In tandem MS, for precursor selection, LMres and HMres were adjusted at 50 V collisional voltage until a single peak was recorded, and then dissociation was induced. To calibrate raw data, CsI (25 mg/ml) spectra were acquired and calibration was carried out with MassLynx (Waters) software. Data were analysed using MassLynx (Waters).

### Small-molecule mass spectrometry

Analytical LC-MS data were obtained using a Waters Alliance 2795-HT LC-MS system equipped with a Phenomenex Kinetex column (2.6 μm, C_18_, 100 A°, 4.6 × 100 mm^2^) and a Phenomenex Security Guard precolumn (Luna, C5, 300 A°) at a flow rate of 1 mL/min at 30 °C. Detection was carried out by a diode array detector (Waters 2998) in the range 210 to 600 nm together with a Waters Quatro-Micro mass detector, operating simultaneously in ES^+^ and ES^−^ modes between 100 and 1000 m/z. Solvents were: A, HPLC grade H_2_O containing 0.05% formic acid; B, HPLC grade CH_3_CN containing 0.045% formic acid. The gradient was as follows: 0 min, 10% B; 10 min, 90% B; 12 min, 90% B; 13 min, 10% B; 15 min, 10% B. Samples (20 µl) were made up in a mixture of buffer and CH_3_CN between 10 and 1000 µg/mL. Enzyme assays were terminated by addition of an equivalent volume of CH_3_CN. Protein was removed by centrifugation and reaction components were directly injected.

## Acknowledgments

We explicitly thank all people who are not included on the author list, in particular those responsible for the PETRA III machine operation, and all other DESY staff who are making this joint effort possible. This research was supported in part through the Maxwell computational resources operated at Deutsches Elektronen-Synchrotron (DESY), Hamburg, Germany, and by the Max Planck Society, the Center for ultrafast imaging and the Joachim Herz Foundation (Biomedical Physics of Infection).

## Supplementary Figures

**Supplementary figure 1:**
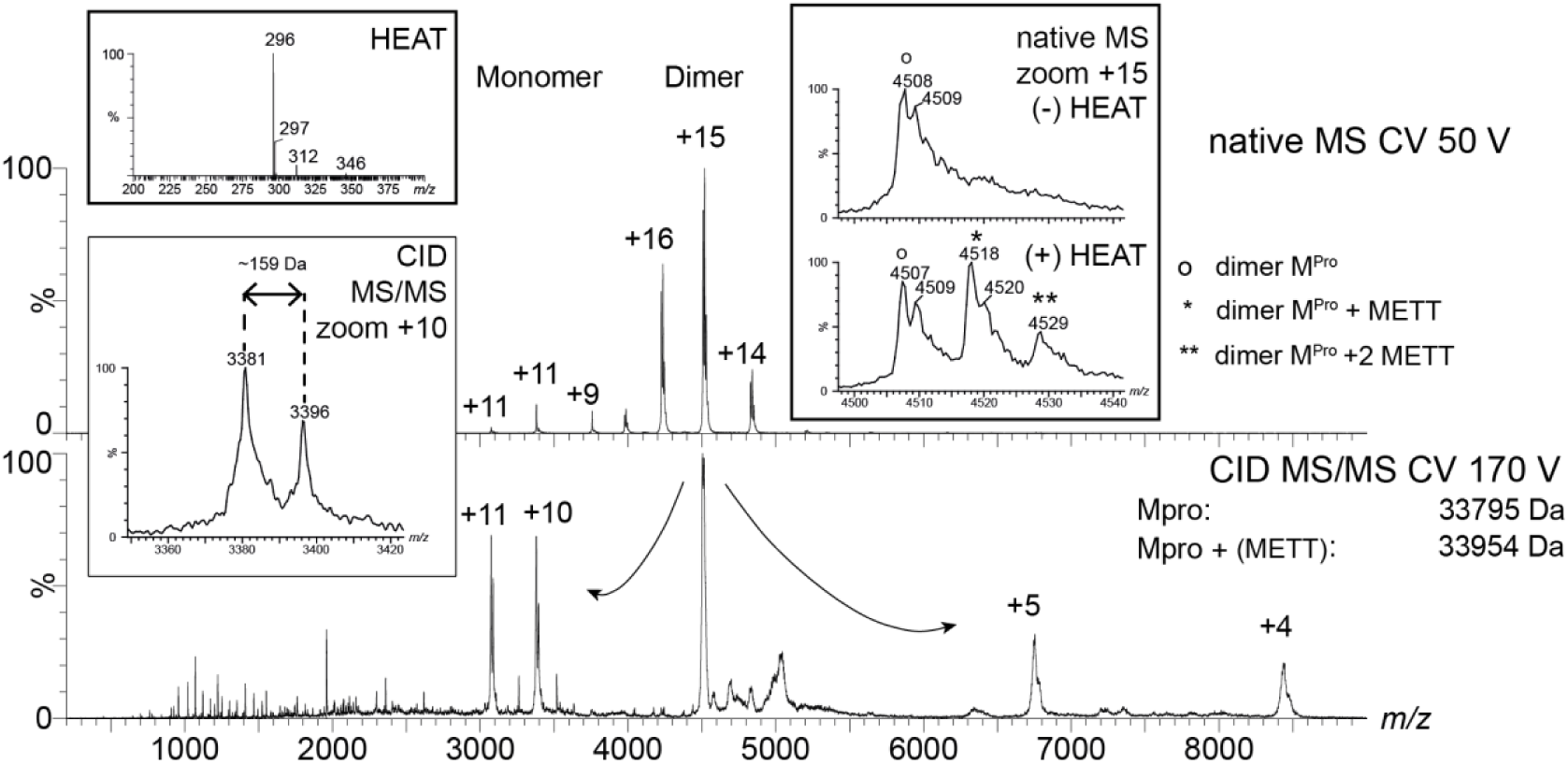
A degradation product of HEAT is covalently bound to M^pro^. Main graphs showing spectra of M^pro^ incubated with HEAT of native MS (top) and after collision induced dissociation (bottom). Insets: top left, intact HEAT after incubation with M^pro^, bottom left, monomeric M^pro^ with and without adduct of METT; right, native MS spectra without (top) and with HEAT (bottom), showing the dimer of M^pro^ as apo and with one or two METT bound.

**Supplementary figure 2:**
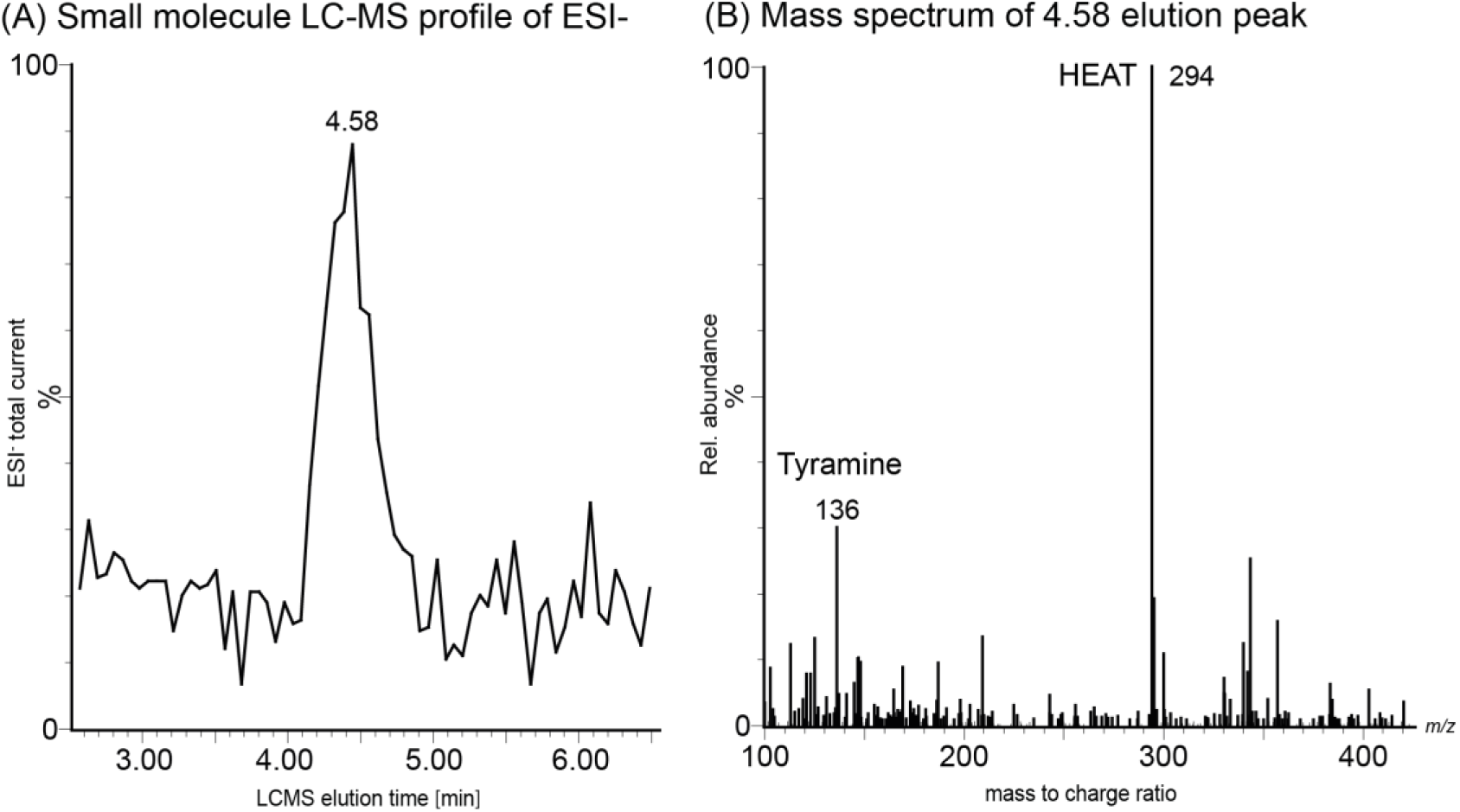
MS analysis of the educts and products of the reaction of HEAT with M^pro^. (A) HPLC elution profile (ESI^-^, Total Ion Current) of the reaction products of HEAT with M^pro^ in the presence of DTT and the (B) corresponding mass spectra with assigned peaks for [TY]^-^ and [HEAT]^-^.

**Supplementary figure 3:**
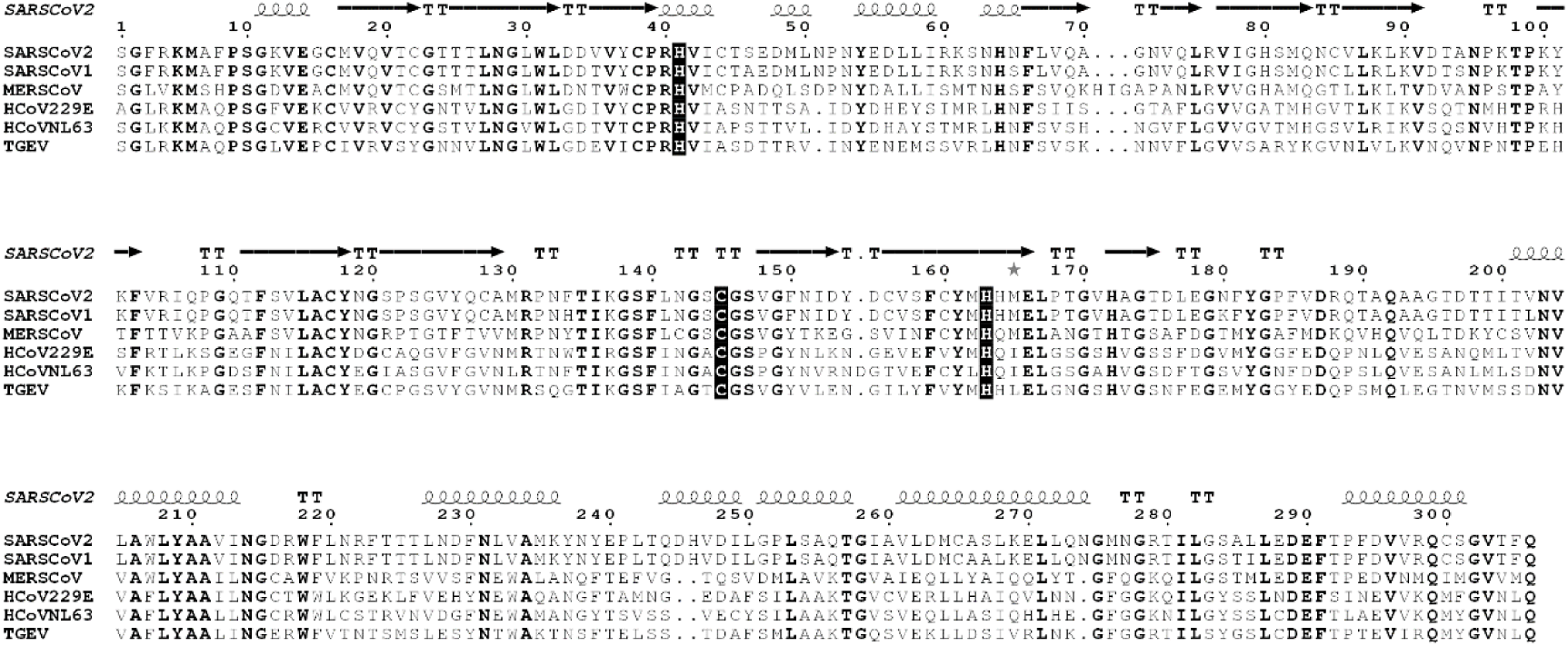
Conservation of M^pro^ active site in α- (HCoV229E, HCovNL63, TGEV) and β- (SARS-CoV-1&2, MERS-CoV) coronaviruses. Conserved residues are depicted in bold. Active site residues His41 and Cys145 are highlighted in black box. Also His163 forming a hydrogen bond to METT is conserved.

